# Investigating Cryopreserved PBMC Functionality in an Antigen-Induced Model of Sarcoidosis Granuloma Formation

**DOI:** 10.1101/2024.03.23.586344

**Authors:** Sarah G. Seman, Sabahattin Bicer, Mark W. Julian, Patrick J. Kramer, Jonah R. Mitchell, Elliott D. Crouser, Landon W. Locke

## Abstract

Sarcoidosis, a systemic inflammatory disease, poses challenges in understanding its etiology and variable clinical courses. Despite ongoing uncertainty about causative agents and genetic predisposition, granuloma formation remains its hallmark feature. To address this complexity, we developed a validated in vitro human granuloma model using peripheral blood mononuclear cells (PBMCs), providing a dynamic platform for studying sarcoidosis pathogenesis. While cryopreservation is a common method for long-term sample preservation, the biological effects of freezing and thawing PBMCs on granuloma formation remain unclear. This study aimed to assess the viability and functionality of cryopreserved sarcoidosis PBMCs within the granuloma model, revealing similar granulomatous responses to fresh cells and highlighting the potential of cryopreserved PBMCs as a valuable tool for studying sarcoidosis and related diseases.

## Introduction

Sarcoidosis is an uncommon heterogeneous systemic inflammatory disease that can affect multiple organs. Understanding of sarcoidosis etiology remains elusive, and questions persist regarding variable clinical courses between patients and the risk for progressive disease. Although described as early as 1877, there is ongoing uncertainty regarding the causative agents of sarcoidosis as well as the predisposing genetic susceptibility leading to active disease. The hallmark histopathological feature of sarcoidosis is the formation of granulomas in target tissues, which have unique characteristics compared to other granuloma-forming diseases ^1^. Efforts to model sarcoidosis and understand its pathogenesis face significant challenges. The lack of a clear link between human gene polymorphisms and sarcoidosis hinders the effective use of animal models, which typically require specific genetic modifications to mimic the disease. While many research teams have introduced animal models, the cellular composition and sustainability of resulting granulomas often do not fully resemble those observed in human sarcoidosis. One exception is the Tsc2 knockout mice created by Weichart *et al*. ^2^, which display tissue granulomas similar to those in sarcoidosis patients with active disease.

We established a simple yet powerful lab model of human sarcoidosis: an *in vitro* human granuloma model ^3,4^. Moving beyond the conventional static examination of cells, our model, utilizing peripheral blood mononuclear cells (PBMCs), facilitates dynamic interactions between antigens and immune cells that are known constituents of sarcoidosis tissue granulomas. The granulomas elicited in the model authentically represent the functional aspects of granuloma formation *in vivo* as demonstrated by the overlap in cellular pathways between patient PBMC-generated granuloma-like structures and diseased sarcoidosis lung and lymph node tissues ^5^. Sarcoidosis granuloma lesions have exhibited macrophages of various subtypes and activation states, with upregulated macrophage genes observed in cardiac sarcoidosis lesions ^6^. Investigation into macrophage activation in sarcoidosis granulomas produced in the model revealed heightened CD163 expression on centrally located macrophages ^3^, mirroring findings in diseased tissues from neuromuscular sarcoidosis ^7^. Thus, the model has the capacity to mimic *in vivo* granulomatous inflammation and can unveil mechanisms of disease pathology.

Cryopreservation, involving the storage of cells below –150°C, serves as a common method for long-term preservation of patient samples, including PBMCs. Incorporating cryopreserved PBMCs into the granuloma model offers several advantages. It would facilitate research collaboration by enabling transportation of cells from various clinics to model laboratories, broadening recruitment networks for specific sarcoidosis phenotypes. Additionally, it would allow for coordinated study of PBMCs from multiple donors, reducing variability in model analysis. However, the biological effects of freezing and thawing PBMCs on granuloma formation are unknown. This study aims to assess viability and functional features of cryopreserved sarcoidosis PBMCs in the granuloma model, including antigen-induced responses and cytokine production.

## Materials and Methods

### Study participants and exclusion criteria

This study was approved by The Ohio State University Institutional Biomedical Sciences Review Board (2014H0380). All study subjects were enrolled in compliance with National Institutes of Health guidelines, and in accordance with the amended Declaration of Helsinki and provided informed written consent. The diagnosis of sarcoidosis conformed to official ATS guidelines ^8^, and we recruited 10 sarcoidosis patients with signs of active organ involvement (**Table 1**). The exclusion criteria included individuals under 18 years old, cigarette smokers, individuals with latent or active tuberculosis (TB) infection, and patients who required treatment with oral corticosteroids and/or immunomodulators within the last 6 months. The Scadding stage for each patient was assessed by a radiologist based on chest X-ray (CXR) imaging.

**Table 1:** Patient demographics. Abbreviations: M, male, F, female, W, White, and B, Black. The Scadding stage is a radiographic classification system derived from chest X-ray findings, used to assess the severity and progression of pulmonary sarcoidosis. Scadding stages range from 0 (no visible abnormalities) to IV (extensive pulmonary fibrosis), as described in reference ^19^.

| Patient # | Age (years) | Gender | Race | Scadding Stage (CXR) | Extra-Pulmonary Disease |
| --- | --- | --- | --- | --- | --- |
| 1 | 44 | M | B | II | Y |
| 2 | 53 | F | B | I | Y |
| 3 | 74 | M | W | I | Y |
| 4 | 64 | F | W | I | N |
| 5 | 53 | M | W | III | N |
| 6 | 56 | M | W | II | Y |
| 7 | 56 | F | B | II | Y |
| 8 | 67 | F | W | II | Y |
| 9 | 57 | F | W | III | N |
| 10 | 53 | M | B | II | Y |

### PBMC isolation and the *in vitro* granuloma model

Patients with sarcoidosis who consented to participate in the study underwent peripheral venipuncture to obtain 120 mL of blood into sodium heparin vacutainers (Becton, Dickinson and Company, Franklin Lakes, NJ). Blood samples were kept at room temperature throughout processing. PBMCs were isolated from whole blood using the Ficoll-Paque method, as previously described ^4^, and adjusted to 2 million cells per mL in RPMI 1640 medium (Gibco, ThermoFisher, Waltham, MA) containing 10% human AB serum. The granuloma model was then initiated by seeding one million PBMCs per well into a 24-well plate. Following a 30-minute equilibration period, 1.0 μm-sized blue fluorescent beads (Polysciences Inc., Warrington, PA) were added to the culture at a target bead load of 50 per monocyte, which is a population assumed to comprise 10% of PBMCs. These beads were either coated with purified protein derivative (PPD, AJ Vaccines, Copenhagen, Denmark) or left uncoated, with triplicate wells prepared for each bead type. Following the challenge, the cells were incubated at 37°C with 5% CO_2_ for 7 days.

### Cryopreservation protocol

Following each isolation performed, approximately half of the PBMCs were cryopreserved. Briefly, PBMCs were suspended in freezing media consisting of 10% dimethyl sulfoxide (DMSO) in heat-inactivated fetal bovine serum (FBS) (FisherBrand; Pittsburgh, PA) at a concentration of 10 million cells/mL. Cell suspensions were aliquoted into cryovials at 1 ml per vial (10x10^6^cells per vial), loaded into a Mr. Frosty freezing container (ThermoFisher, Waltham, MA), and immediately placed in a –80°C freezer. After 24 hours, the cryovials were transferred to a liquid nitrogen storage tank (ThermoFisher, Pittsburgh, PA) maintained at –196°C for long-term preservation.

### Thawing procedure

Thawing media, consisting of RPMI 1640 medium with 10% FBS, was freshly prepared and warmed to 37°C. Cryovials containing PBMCs were retrieved from the liquid nitrogen storage tank and partially immersed in a 37°C water bath for 2 minutes to initiate controlled thawing. The cell suspension from each cryovial was carefully transferred to an excess of pre-warmed thawing media in 15 mL conical tubes. Subsequently, the PBMCs underwent centrifugation at 300 x g for 6 minutes at room temperature. After discarding the supernatant, the cell pellet was resuspended in RPMI medium. The cell concentration was determined, and the suspension was adjusted to a density of 2 million cells per mL in RPMI medium supplemented with 10% AB human serum. Following these steps, the granuloma model was initiated as previously described ^4^.

### Granuloma imaging and area fraction quantification

Imaging of the granuloma-like structures and cells was performed on day 7. Briefly, the well plate was placed into a stage-top incubation chamber (Okolab Inc., Sewickley, PA) to maintain the cells at 5% CO_2_ and 37°C. The stage-top chamber was set onto the stage of a Stellaris 5 confocal microscope (Leica Microsystems, Deerfield, IL). Transmitted (brightfield) and blue fluorescence images of the granuloma-like structures and cells were acquired via confocal laser scanning microscopy (CLSM) using a 10x Plan Apo dry objective lens (NA 0.40). Blue bead excitation was facilitated by a 405 nm diode laser, with the emission range set to 410 nm to 530 nm. Image resolution was set to 1024x1024 pixels, and no averaging was applied. At least five images from three different wells were acquired for both uncoated and PPD bead-challenged PBMCs. Images were acquired at random locations within each well, avoiding the direct center due to locally higher cell density that obstructs granuloma detection. The images were imported and analyzed using a semi-automated analysis script in Materials Image Processing and Automated Reconstruction (MIPAR™ v2.2.5; Worthington, OH) software, following methodology previously described ^3^. The degree of granuloma formation was quantified as the percentage of the area occupied by granuloma-like structures relative to the total image field of view.

For select donors, we assessed the relative fractions of viable cells compared to dead cells at day 7 in the granuloma-like structures. To visualize live cells apart from dead ones, the calcein AM and ethidium homodimer dyes were added to the cells at final concentrations of 2 μM and 4 μM, respectively. After a 30-minute incubation, the well plates were loaded into the Okolab chamber for imaging while being maintained at 5% CO_2_ and 37°C. Using a 40x water immersion objective, three-dimensional z-stack images were acquired at a resolution of 2048x2048 pixels, corresponding to an X-Y pixel size of 400 nm by 400 nm. The 405 nm, 488 nm, and 548 nm lasers were used to image the blue fluorescent beads, calcein AM, and ethidium homodimer, respectively. To minimize signal cross-talk, images were acquired in line sequential mode and no line averaging was applied. Single stain control samples were prepared to confirm no signal crosstalk between the emission channels.

### Supernatant collection and Multiplex ELISA

Cell culture supernatants from the *in vitro* cultures were collected on day 7. Briefly, the supernatants from triplicate wells were collected and immediately centrifuged at 200 x g for 10 minutes to remove cellular debris. The clarified supernatants were stored at –80°C until analysis. Cytokine levels in the culture supernatants were measured using an MSD^®^ Multi-Spot Assay System, V-Plex Proinflammatory Panel 1 human Kit (K15049D-1, MesoScale Discovery LLC, Maryland, U.S.A), following the manufacturer’s instructions. The V-Plex plate was washed three times prior to the addition of analyte standards and 1:2 diluted sample supernatants. After another round of washing, the detection antibodies were added to the wells. Following a final wash step, MSD read buffer was added to each well, and signals were measured using an MSD Quickplex SQ120 instrument. Analyte standards and samples were run in duplicate and averaged. The Discovery Workbench software was used to generate a calibration curve for each analyte and to calculate the analyte concentration in pg/mL. Concentration values surpassing the upper limit of detection (ULOD) were reassigned the ULOD value determined from the standard curve for each analyte.

### Statistical Analysis

Statistical analyses were performed using GraphPad Prism 10.0 software (GraphPad Software, Boston, MA). A paired t-test was employed to assess differences in cytokine production and granuloma area fraction between fresh and cryopreserved PBMCs. Linear regression analysis was utilized to examine the relationship between cryopreservation storage duration and the recovery of viable PBMCs. Group values are mean ± SEM; a *p*-value less than 0.05 was considered statistically significant.

## Results

### Demographic and clinical characteristics of enrolled patients

All 10 patients enrolled into the study were diagnosed with biopsy-confirmed pulmonary sarcoidosis (**Table 1**), with seven of the patients exhibiting evidence of disease outside of the lungs. Their mean age was 58 years ± 9 (SEM) (range: 44–74), with an equal distribution of male and female patients. Among the cohort, 6 were of white ethnicity, while 4 were black. The CXR-based Scadding stage assigned to each patient ranged from I to III, with no patients assigned a stage of 0 or IV.

### PBMC viability is preserved over extended cryopreservation storage durations

The duration of PBMC cryopreservation storage ranged from 7 to 84 days. Linear regression analysis revealed no statistical correlation between cryopreservation storage time and recovery (**Figure 1**). Notably, additional samples from our laboratory exhibited a consistent recovery rate exceeding 60% even after 450 days of cryopreservation storage (data not shown).

**Figure 1.**
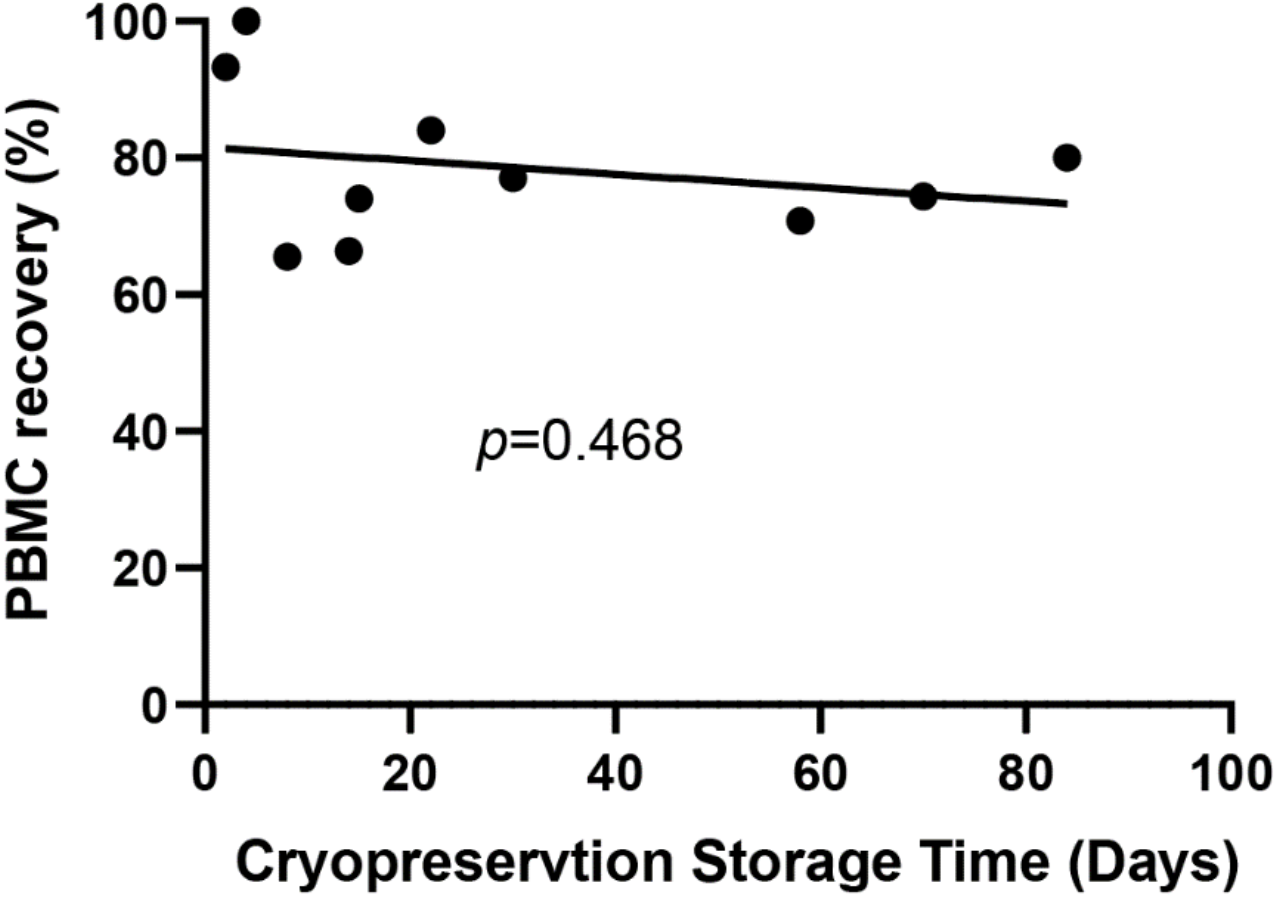
PBMC viability is preserved over cryopreservation storage durations. Percentage of PBMC recovery post-thaw from liquid nitrogen storage. Linear regression analysis revealed no statistically significant correlation between storage time and cell recovery (*p*=0.468).

### Cryopreserved sarcoidosis PBMCs maintain granuloma responses in the *in vitro* model

We first aimed to determine whether cryopreserved PBMCs retain similar granulomatous responses as fresh cells to bead stimulations. Upon incubation of freshly isolated sarcoidosis PBMCs with uncoated beads at day 7, we observed minimal cell aggregation and a correspondingly low granuloma area fraction, aligning with our prior observations ^3^. In contrast, when these PBMCs were incubated with PPD coated beads over 7 days, a robust granuloma response was triggered in 6 out of 10 patients. Here, a high response was defined as a granuloma area fraction greater than or equal to 2%. Four patients exhibited a low granulomatous response (area fraction less than 2%) to PPD bead stimulation.

In the analysis of patient-matched cryopreserved PBMCs, the computed granuloma area fraction for uncoated bead stimulation remained minimal and exhibited no statistically significant difference from the fresh response, (**Figure 2A)**. Likewise, the granulomatous response of cryopreserved cells to PPD coated beads showed no statistically significant difference compared to fresh cells (**Figure 2B**). **Figure 2C** shows images illustrating the overlaid color-coated granuloma regions for a representative patient.

**Figure 2:**
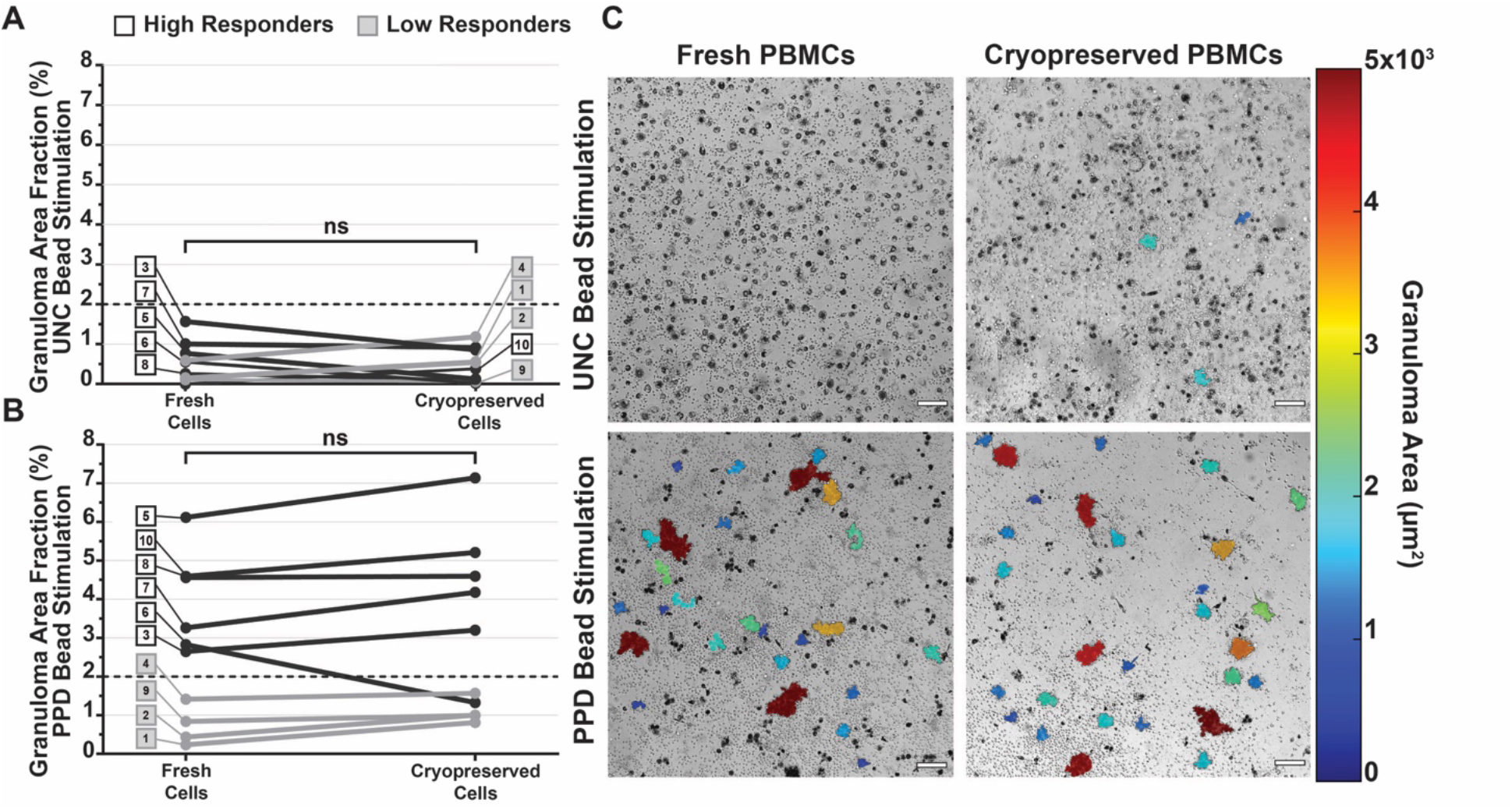
Image analysis demonstrates that granuloma formation is preserved in the *in vitro* model using cryopreserved PBMCs from sarcoidosis patients. (**A**) The calculated granuloma area fraction following uncoated bead stimulation at day 7 showing no statistically significant difference between patient-matched fresh and cryopreserved cells. (**B**) Granuloma area fraction in response to purified protein derivative (PPD) coated beads at day 7 did not exhibit a statistically significant difference between fresh and cryopreserved cells (**C**) Representative images with color overlay of detected granulomas at day 7 post-stimulation with uncoated (top row) and PPD coated beads (bottom row) for freshly isolated and cryopreserved cells from the same patient.

### Granuloma-like structures derived from cryopreserved PBMCs feature high cellular viability

Having established that cryopreserved PBMCs maintained their ability to form granuloma-like structures in response to PPD bead stimulation, we next assessed the viability of these structures at the day 7 timepoint. CLSM 3D-rendered z-stack images, acquired following the addition and incubation of calcein AM and ethidium homodimer (live and dead cell indicator dyes, respectively), revealed that the majority of cells comprising the granuloma-like structures were viable. This was observed for both uncoated bead-stimulated cells (**Figure 3A-D**) and PPD bead-stimulated cells (**Figure 3E-H**). Notably, only a minority of cells within and around the granulomas were identified as dead or undergoing cell death processes.

**Figure 3:**
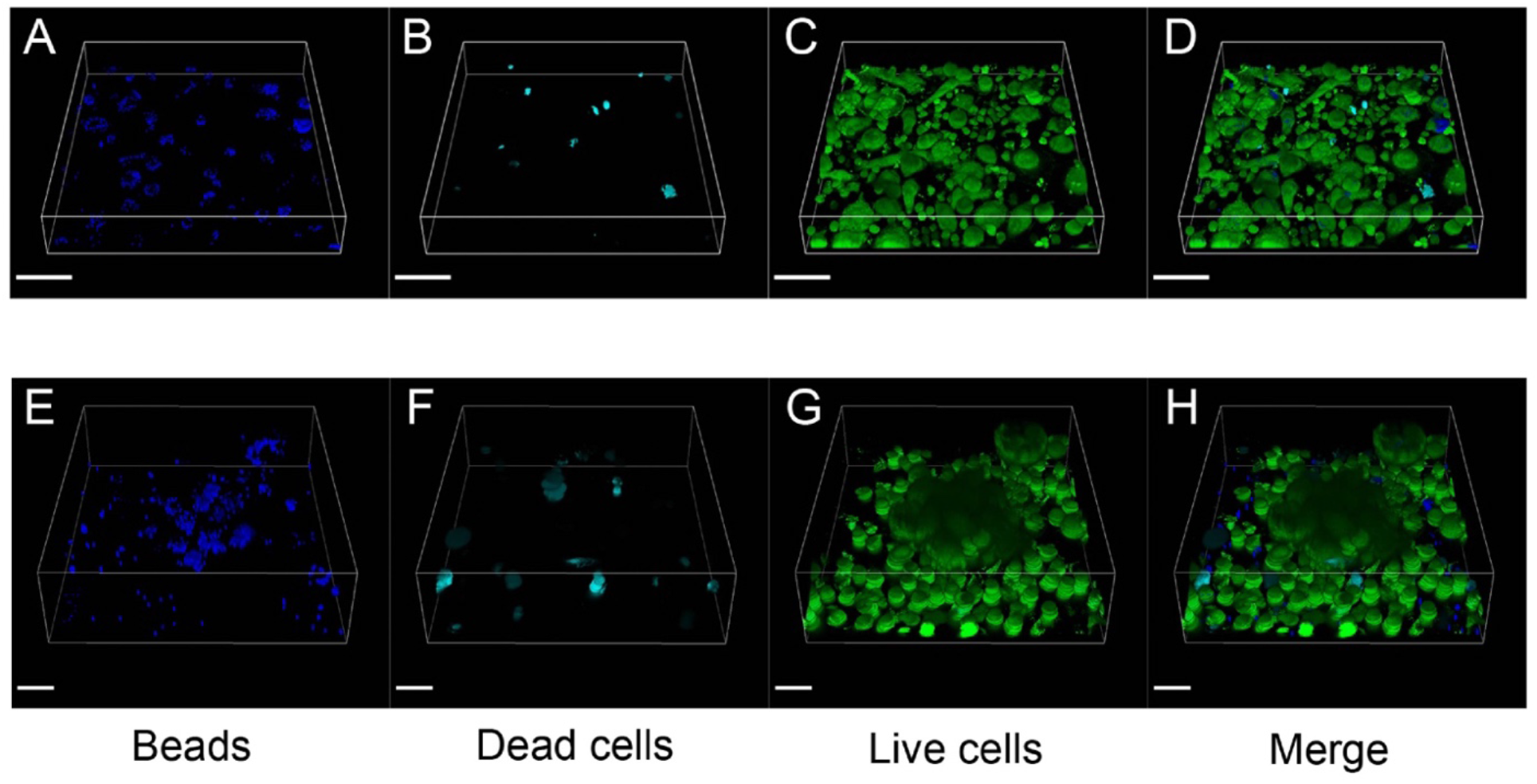
Granuloma-like structures derived from cryopreserved PBMCs feature high cellular viability. Representative confocal laser scanning microscopy images at day 7 post-stimulation with (**A-D**) uncoated beads or (**E-H**) purified protein derivative (PPD) coated beads. Individual channels show (**A, E**) blue fluorescent beads, (**B, F**) ethidium homodimer staining for dead cells (red), and (**C, G**) calcein AM staining for live cells (green). Merged images (**D, H**) demonstrate the predominance of viable cells within the granuloma-like structures. Scale bar: 100 μm.

### Cryopreserved PBMCs retain similar cytokine responses to freshly isolated cells

We have previously shown in the *in vitro* human granuloma model that sarcoidosis granulomas produce a unique cytokine signature compared to other groups ^4^. To examine whether cryopreserved cells, when stimulated with uncoated and PPD coated beads, maintain similar cytokine production patterns as observed in freshly isolated cells, we chose to examine 10 cytokines representing both pro- and anti-inflammatory cytokines. For each cytokine analyzed, we present the ratio of its measured concentration between PPD and uncoated bead stimulation for both fresh and cryopreserved cells. We found that for most cytokines, namely IL-2, IL-4, IL-10, IL-12, IL-1β, and IFN-γ, this ratio exhibited statistically non-significant differences between fresh and cryopreserved PBMCs (**Figure 4**). Only two cytokines, IL-13 and TNF-α, demonstrated statistically significant differences between fresh and cryopreserved PBMCs, with cryopreserved cells displaying a reduced ratio. Two cytokines, IL-6 and IL-8, consistently showed measured concentration values that surpassed the upper limit of detection (ULOD), prompting us to omit these cytokines from subsequent analysis.

**Figure 4:**
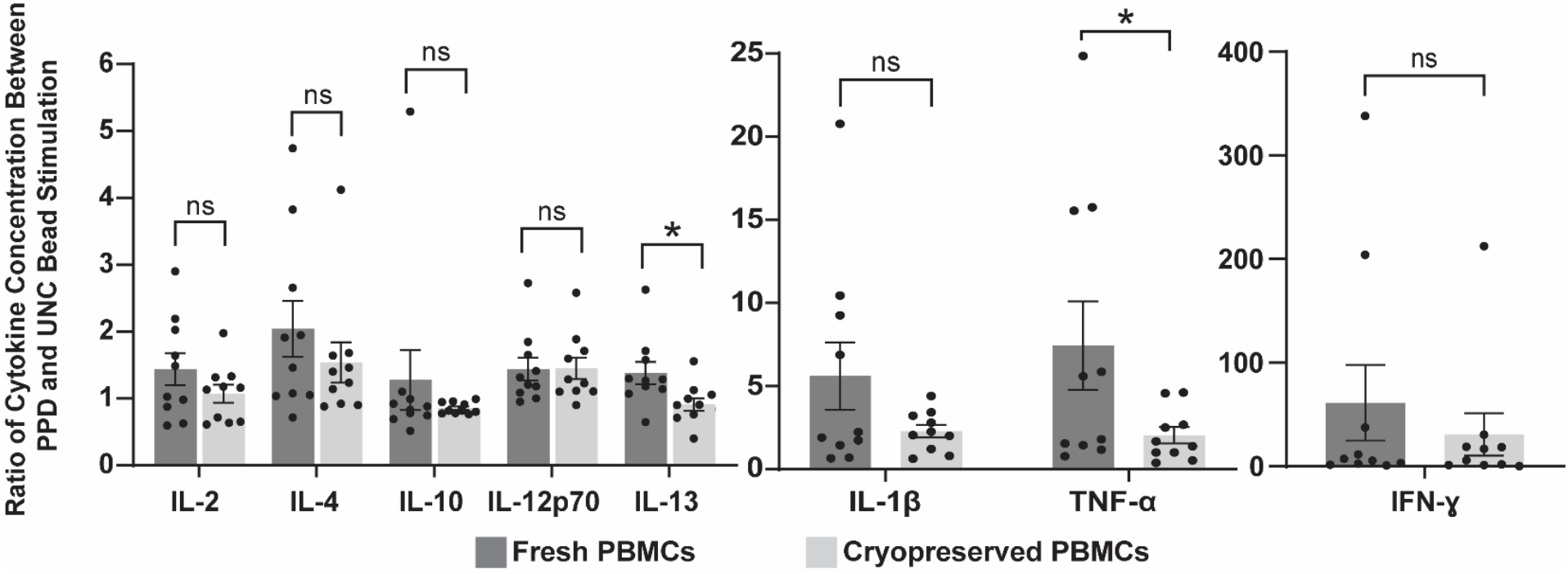
Cryopreserved PBMCs retain similar cytokine responses to freshly isolated cells in the granuloma model. Cytokine production ratios between PPD and uncoated bead stimulations (PPD/uncoated) in fresh and cryopreserved sarcoidosis PBMCs at day 7. Cytokines are grouped based on concentration similarities for visual clarity. ^*^ indicates *p* < 0.05 and ns indicates not significant.

## Discussion

The *in vitro* granuloma model based on patient-derived PBMCs has provided valuable insights into the pathobiology of sarcoidosis. The use of cryopreserved PBMCs from sarcoidosis patients offers several advantages, yet the biological effects of freezing and thawing PBMCs on granuloma formation are unknown. PBMCs contain multiple immune cell types, and differences in the number and viability may be expected among the populations following cryopreservation. Although decreases in cell count have been reported post-cryopreservation, other studies have reported that percentages of specific T cell subsets and regulatory T cells are consistently maintained up to 12 months after cryopreservation ^9^. Importantly, this mode of storage has been shown to maintain specific cell markers and functional responses compared to measurements made on fresh cells ^10^. Previous studies have demonstrated that cryopreserved PBMCs retain key inflammatory responses when compared to freshly isolated PBMCs ^11,12^. For example, Solati *et al*. showed that cryopreserved human monocytes, in response to LPS, exhibit cytokine production levels comparable to donor-matched fresh cells ^13^. Similarly, Kilani et al. ^14^ and Kreher et al. ^15^ found that cytokine production of cryopreserved lymphocytes did not deviate from freshly isolated cells.

There have been no prior studies that have demonstrated that cryopreserved PBMCs derived from a rare disease patient group such as sarcoidosis retain similar granulomatous responses in functional cell models. We first established that sarcoidosis PBMCs cryopreserved in 10% DMSO in FBS exhibited minimal reductions in both cell count and viability. All patient samples had less than 35% cell loss during long term storage (up to 84 days of storage). This result indicates the feasibility of cryopreserving sarcoidosis PBMCs for the purposes of transportation as well as when batched experimentation would be preferable.

We next evaluated functional cellular responses in the *in vitro* granuloma model using PBMCs from sarcoidosis patients, comparing fresh and cryopreserved cells. Our analysis centered on assessing granuloma formation in response to both uncoated and PPD coated beads at the day 7 timepoint, leveraging quantitative image data analysis techniques. Our findings revealed no significant disparities in the degree of granuloma formation between fresh cells and cryopreserved cells. Furthermore, utilizing live/dead discriminating dyes, our analysis showed that the majority of cells associated with granulomas remained viable at day 7 in both cryopreserved and fresh cells. Approximately 20% of cells were identified as dead or in the process of dying, a level consistently observed in the model at this timepoint. This outcome demonstrates the compatibility of cell cryopreservation with an *in vitro* granuloma model that features multicellular granulomatous responses.

Besides analyzing the physical formation of granulomas, we also quantified cytokine responses. Cytokines play a pivotal role in regulating immune responses during granulomatous inflammation ^16^ and we previously highlighted distinctive extracellular cytokine patterns in sarcoidosis PBMCs compared to individuals with latent TB infection and healthy controls ^17^. Among the cytokines examined, statistical analysis conducted on the ratio of measured concentrations between PPD and uncoated bead stimulation revealed no significant differences between fresh and cryopreserved responses for 6 out of the 8 tested. However, IL-13 and TNF-α displayed statistical significance, showing a diminished ratio of PPD to uncoated bead stimulation in cryopreserved cells compared to freshly isolated cells.

The observation regarding TNF-α is particularly interesting, where the reduction in ratio stemmed from elevated responses to uncoated beads in cryopreserved cells, rather than diminished responses to PPD coated beads. Interestingly, the increased uncoated bead response observed by cryopreserved cells did not lead to an increased granuloma response. In fact, the granuloma response to uncoated beads by cryopreserved cells was slightly below that of fresh cells, though not statistically significant.

Our study is subject to several limitations. Due to constraints in patient cell availability, we did not examine potential losses within distinct cell populations resulting from the cryopreservation or thawing processes. However, if losses due to cryopreservation did occur in distinct cell populations, they did not translate to significant alterations in granuloma formation in response to antigenic bead challenge. Additionally, we did not test different anti-coagulants at the time of blood collection. While heparin was the anti-coagulant used exclusively in this study, the choice of anticoagulant can influence the expression of surface markers on immune cells, affecting their functional responses. Stone *et al*. ^18^ found that when blood was collected in heparin, followed by PBMC isolation, the expression of CD14 (a co-receptor for bacterial lipopolysaccharide) on monocytes was higher compared to other anticoagulants. This higher CD14 expression suggests enhanced responsiveness to inflammatory stimuli in heparinized samples. Thus, the anticoagulant used during blood collection can impact immune cell activation status and functional responses, which should be considered when interpreting results.

In conclusion, our study demonstrates the feasibility and utility of cryopreserving PBMCs from sarcoidosis patients for *in vitro* modeling purposes, allowing for the investigation of granuloma formation and cytokine responses. Despite potential variations in cell recovery and viability post-cryopreservation, our findings indicate minimal alterations in granuloma formation between fresh and cryopreserved cells. Moreover, cytokine responses remain largely consistent, with only slight disparities observed in select cytokines. While our study has its limitations, including the lack of examination into potential losses within distinct cell populations, it highlights the compatibility of cryopreserved PBMCs with *in vitro* functional granuloma models, offering a valuable tool for studying the pathobiology of sarcoidosis and potentially other granulomatous diseases.

## Acknowledgements

The authors wish to thank Harrison Lee, M.S., C.C.R.P. (The Internal Medicine/Pulmonary, Critical Care and Sleep Clinical Trials Office, Columbus, OH) for his assistance in recruiting and consenting patients for their participation in the IRB-approved experimental study.

## CRediT authorship contribution statement

**Sarah Seman:** Investigation, Writing – review & editing

**Sabahattin Bicer:** Investigation, Conceptualization

**Mark Julian:** Investigation, Writing – review & editing

**Patrick Kramer:** Investigation, Writing – review & editing

**Jonah Mitchell:** Investigation

**Elliott Crouser:** Conceptualization, Writing – review & editing, Supervision, Funding acquisition

**Landon W. Locke:** Project administration, Supervision, Writing – original draft, Conceptualization, Funding acquisition

